## Supporting information for "Release of lipid nanodiscs from charged nano-droplets in the electrospray ionization process revealed by microsecond atomistic molecular dynamics simulations"

**Table of contents**

S1. Supplementary methods

S1.1 Treatment of nonbonded interactions.

S2. Supplementary tables

S3. Supplementary figures

### S1. Supplementary methods

S1.1 Treatment of nonbonded interactions.

The non-bonded interactions in molecular force field mainly include van der Waals and electrostatic interactions, which are usually calculated with the Lennard-Jones (LJ) potential energy function, and the Coulomb potential energy function respectively.

$$U_{LJ}(r) = 4\epsilon_{ij}\left[\left(\frac{\sigma_{ij}}{r}\right)^{12} - \left(\frac{\sigma_{ij}}{r}\right)^{6}\right]$$

$$U_{el}(r) = \frac{q_i q_j}{4\pi\epsilon_0\epsilon_r r}$$

The Van der Waals interaction decays rapidly with the increase of distance ($r$), but the electrostatic interaction is proportional to $r^{-1}$, and the attenuation speed with the increase of distance is much slower than that of van der Waals interaction. So the calculation of electrostatic interactions is the most time-consuming in molecular dynamics (MD) simulations.

Two protocols to calculate non-bonded interactions were tested for our sytems (Table S1). Fistly, we tried to calculate non-bonded interactions without truncation; this strategy has been adopted in reported MD simulations of the ESI processes with thousands to tens of thousands of atoms [1-4] . However, the calculation speed is not feasible for our systems with more than 250,000 atoms. Then the long-range electrostatic interactions were omitted to test the calculation speed. The nonbonded interactions were truncated at either 12/33/67 Å and shifted from 9/30/64 Å, respectively. The calculation speed with a cutoff of 67 Å is almost that of the second protocol, in which the equilibrated systems were located at the center of a huge cubic vacuum box of 1000 Å×1000 Å×1000 Å (Figure 1b), which was large enough to avoid interactions between adjacent images when evaporation occurs. And the nonbonded interactions were truncated at 12 Å with a shift at 10 Å. The Particle Mesh Ewald (PME) [5] method was used to calculate the long-range electrostatic interactions. To speed up the simulations, the box size was reduced to 500 Å×500 Å×500 Å when less than two water molecules were evaporated in each run. The calculation speeds for different treatment of nonbonded interactions were listed in Table S2.

**S2. Supplementary tables**

Table S1. Overview of the simulations performed in this study.

| System net charge | Non-bonded interactions | Temperature (K) | Replicas of simulations | Simulation time[a] |
|---|---|---|---|---|
| +69 | PME, Cutoff = 12 Å | 370→450 | 3 (450-1, 450-2 and 450-3) | 150 ns |
| +69 | PME, Cutoff = 12 Å | 370 | 3 (370-1, 370-2 and 370-3) | 450 ns |
| +69 | PME, Cutoff = 12 Å | 300 | 4 (300-1, 300-2, 300-3 and 300-4) | 1200 ns |
| +69 | Cutoff = 12 Å | 370→450 | 1 (C12) | 150 ns |
| +69 | Cutoff = 33 Å | 370→450 | 1 (C33) | 150 ns |
| 0 | PME, Cutoff = 12 Å | 370→450 | 1 (Neutral) | 150 ns |

[a]: simulation time for each trajectory.

Table S2. Simulation performance speeds for different methods of non-bonded interaction calculations (720 CPUs).

| | Cutoff (Å) | Number of atoms | Periodic cubic box size (Å) | Speed (days/ns) |
|---|---|---|---|---|
| Cutoff | 12 | 250,609 | - | 0.23 |
| Cutoff | 33 | 250,609 | - | 0.84 |
| Cutoff + PME | 12 | 250,609 | 1000 | 1.63 |
| Cutoff | 12 | 151,237 | - | 0.16 |
| Cutoff | 33 | 151,716 | - | 0.48 |
| Cutoff + PME | 12 | 150,105 | 1000 | 1.56 |
| Cutoff | 12 | 30,418 | - | 0.06 |
| Cutoff | 33 | 28,927 | - | 0.29 |
| Cutoff + PME | 12 | 27,972 | 500 | 1.03 |

**S3. Supplementary figures**

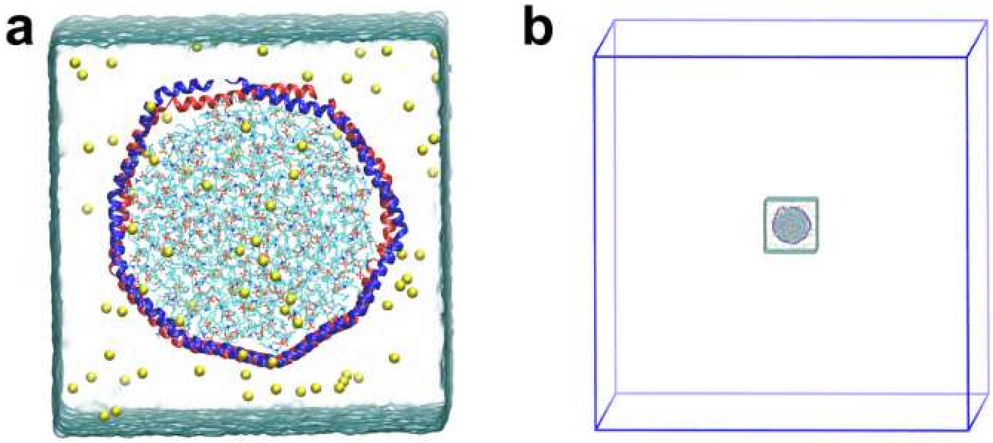


Figure S1. Setup of the nanodisc systems. (a) A close-up of the system. The system contains a MSP1 dimer (charge -12) with two monomers colored in blue and red respectively, 186 DMPC lipids, and 81 sodium ions (yellow spheres). The water is shown in glass bubble surface. (b) The system for the simulations of the ESI process. The periodic boundary box is shown in blue solid lines.

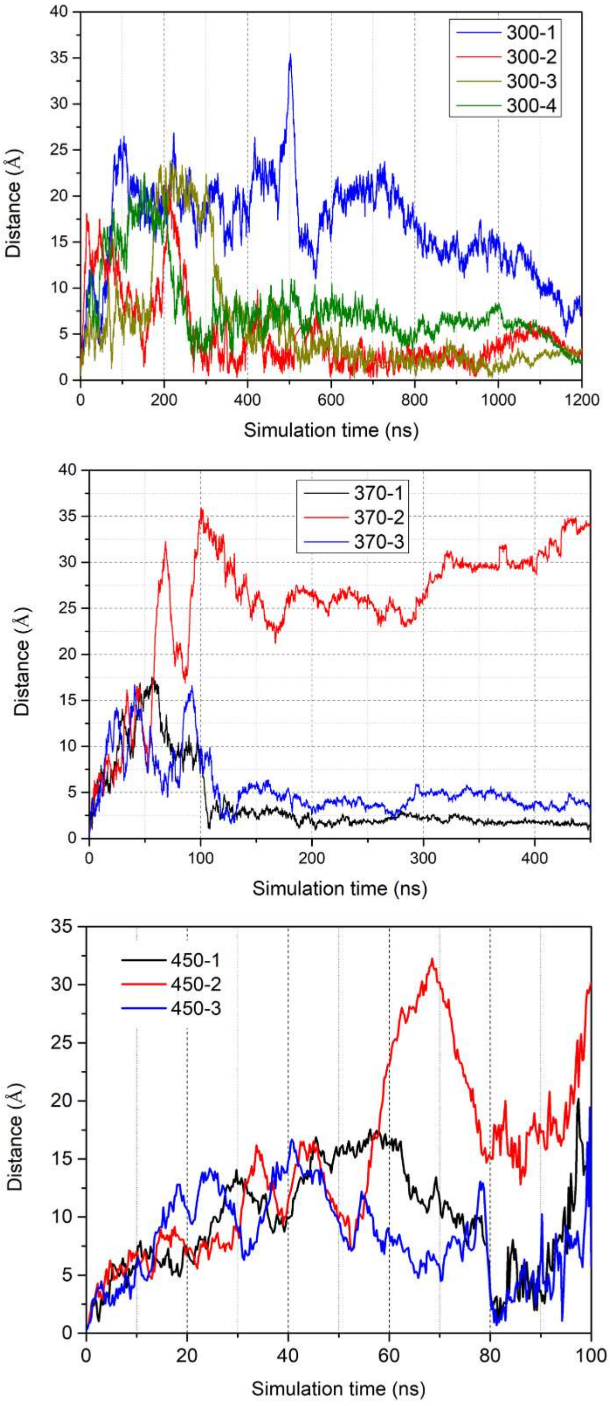


Figure S2. The variation of distances between centers of mass of water molecules and the nanodisc over simulation times.

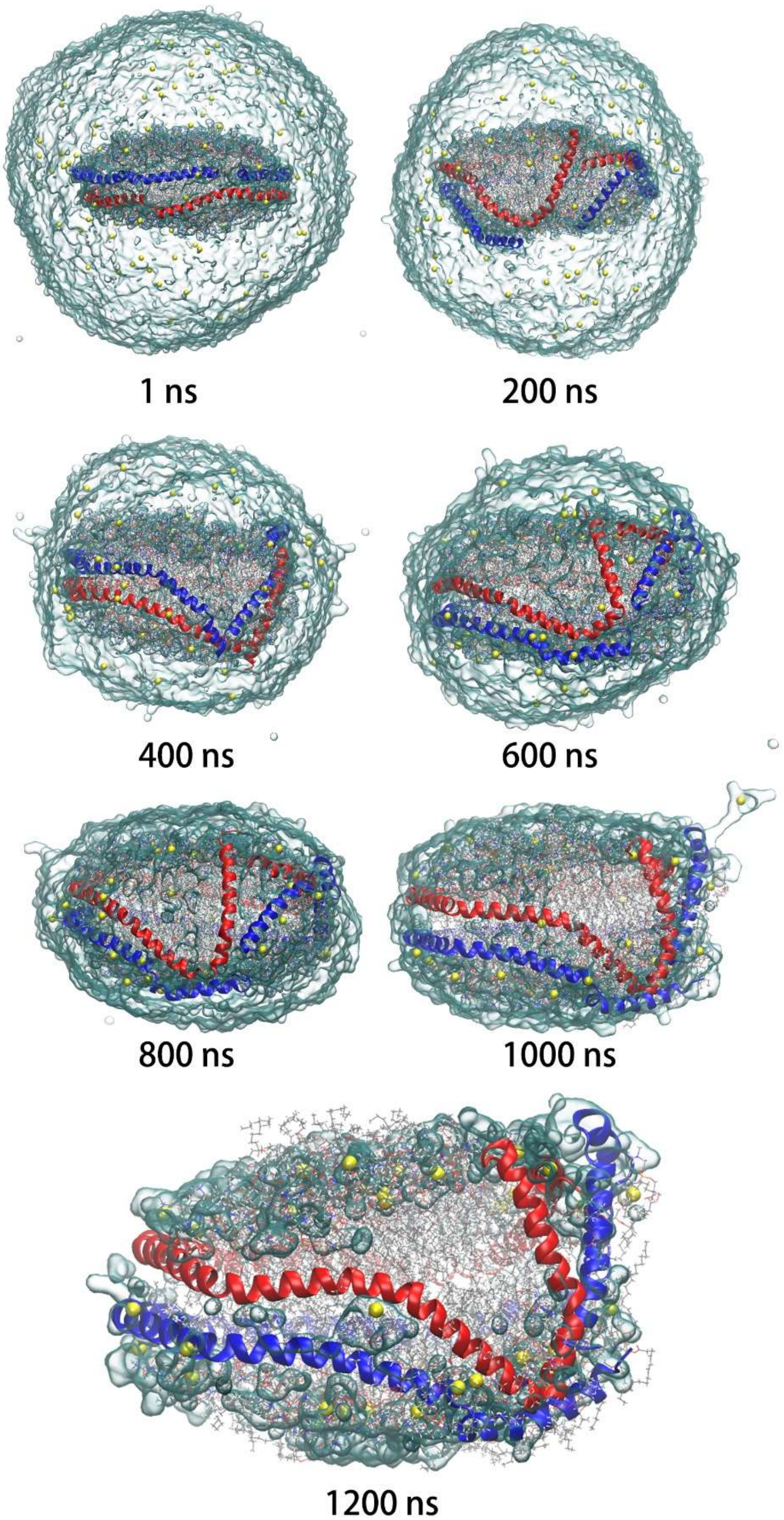


Figure S3. Evolution of the charged nanodisc droplet in the 300-2 simulation. The representation is the same as Figure 1.

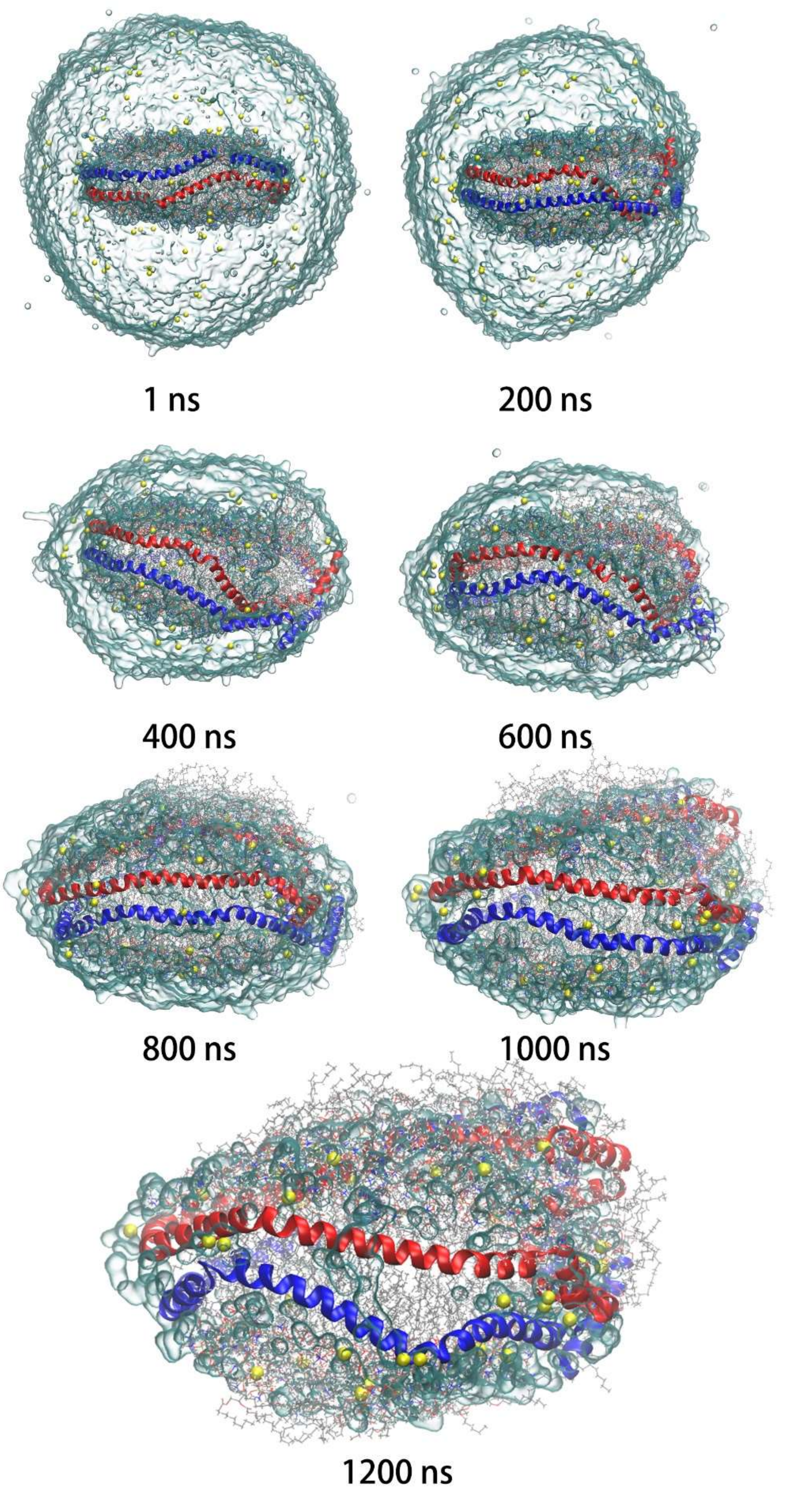


Figure S4. Evolution of the charged nanodisc droplet in the 300-4 simulation. The representation is the same as Figure 1.

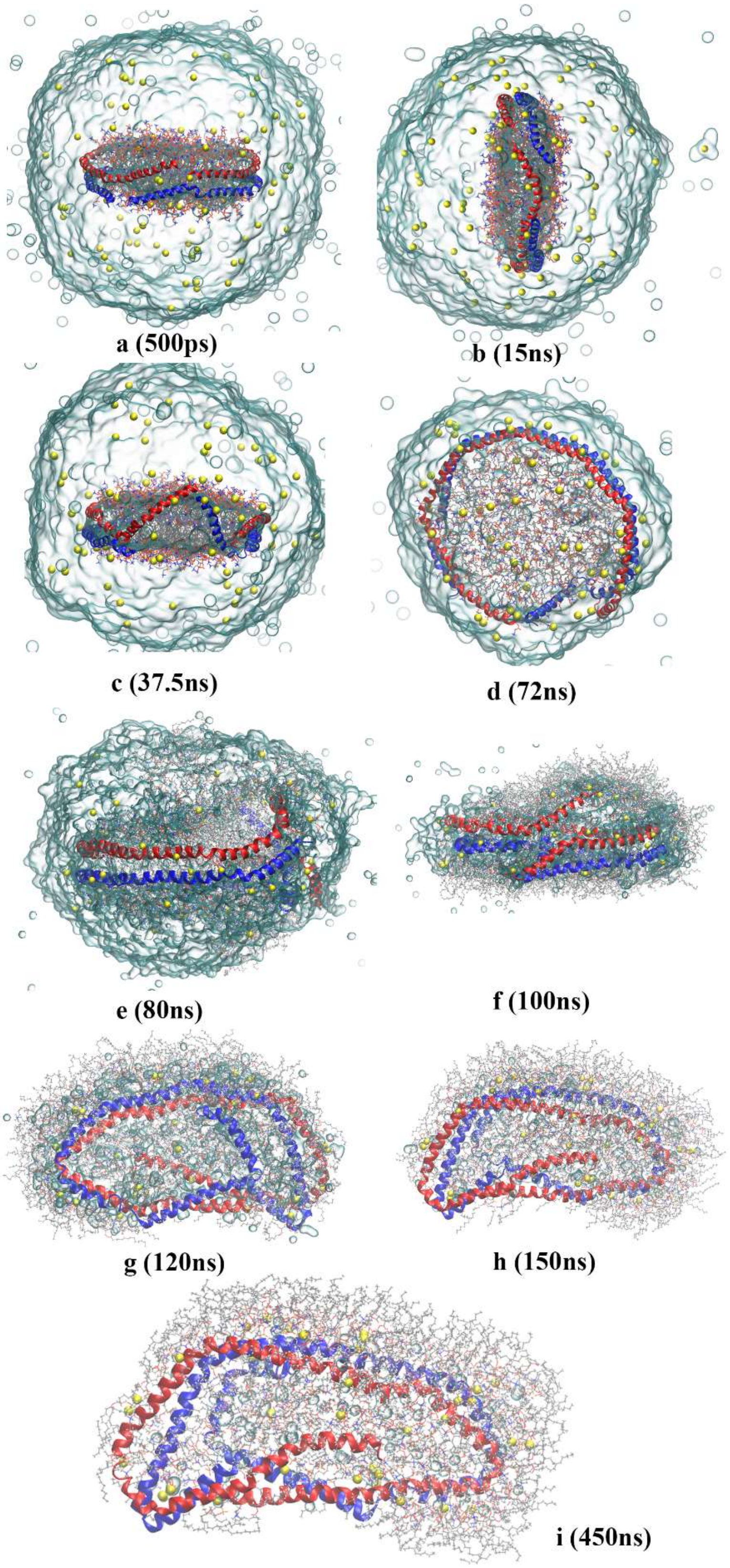


Figure S5. Evolution of the charged nanodisc droplet in the 370-1 simulation. The representation is the same as Figure 1.

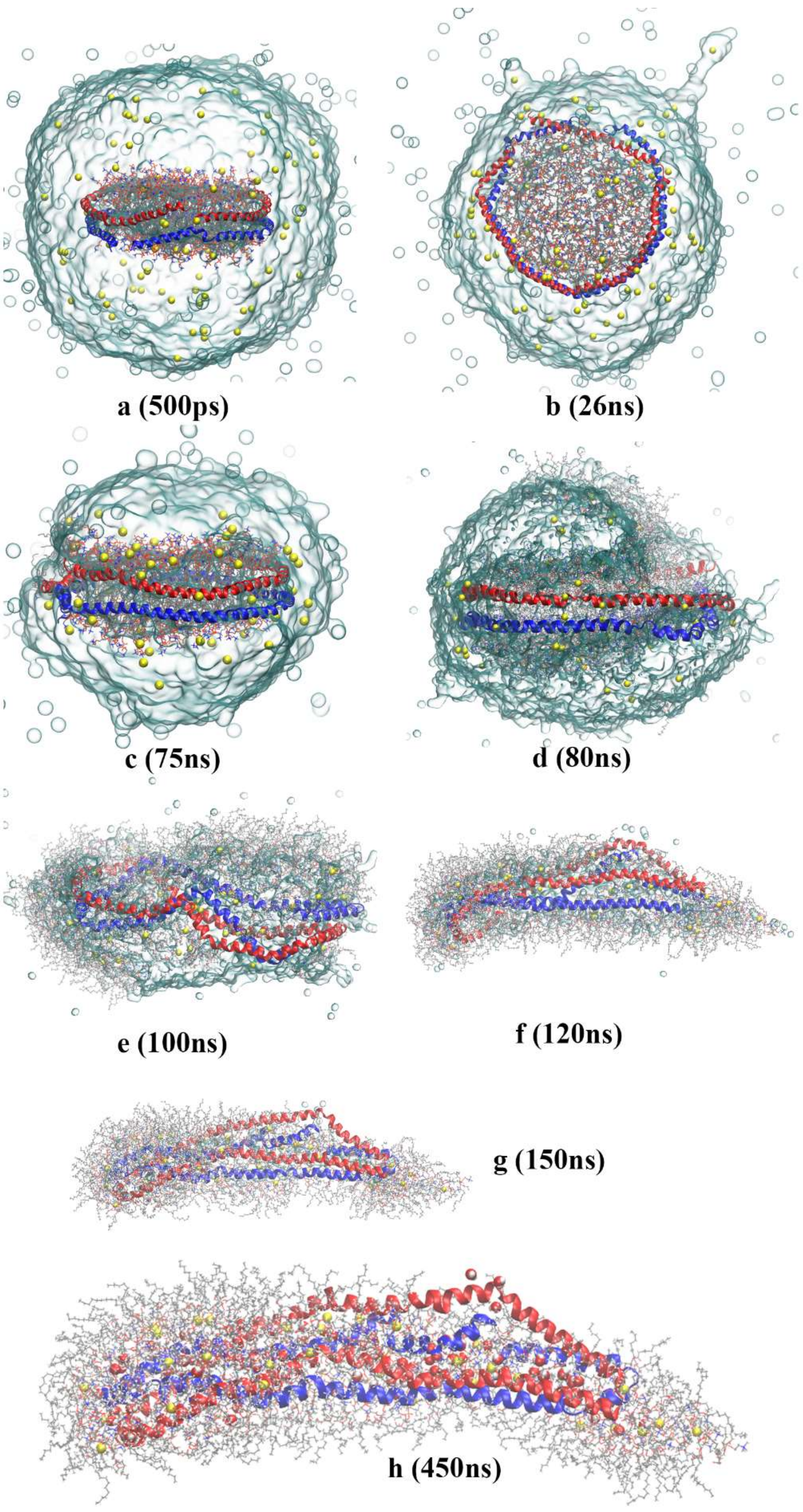


Figure S6. Evolution of the charged nanodisc droplet in the 370-3 simulation. The representation is the same as Figure 1.

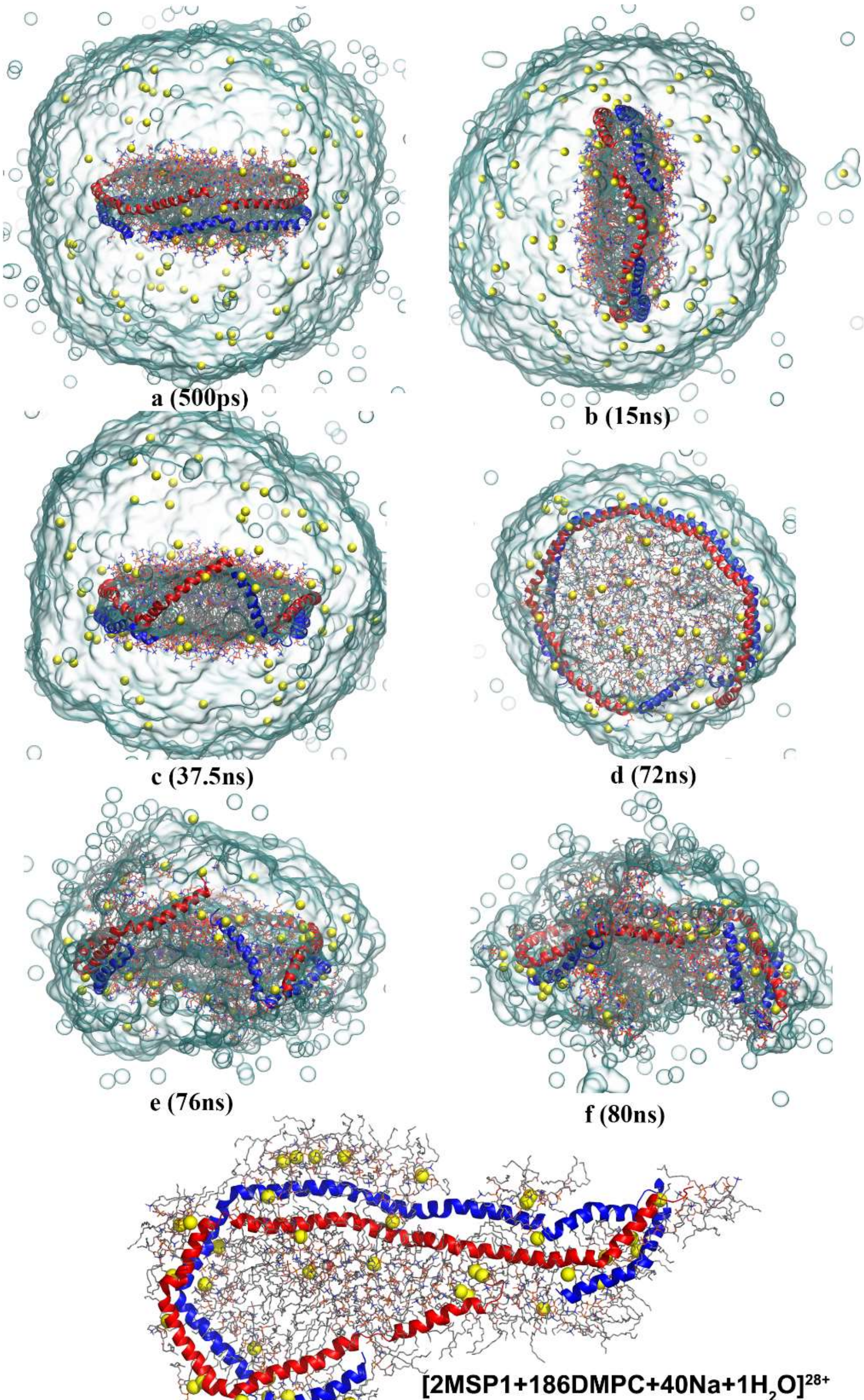


Figure S7. Evolution of the nanodisc droplet in the 450-1 simulation. The representation is the same as Figure 1.

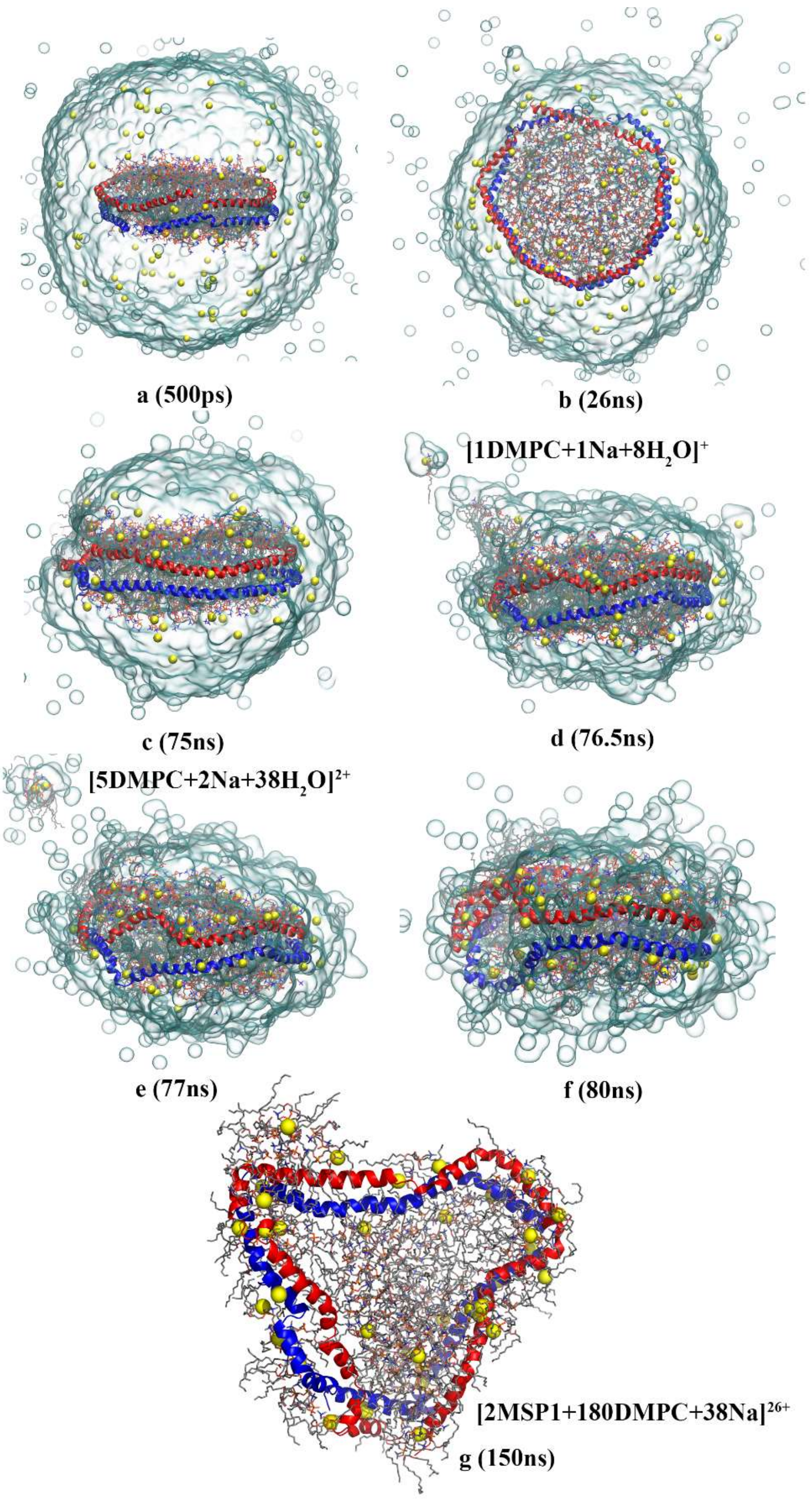


Figure S8. Evolution of the charged nanodisc droplet in the 450-3 simulation. The representation is the same as Figure 1.

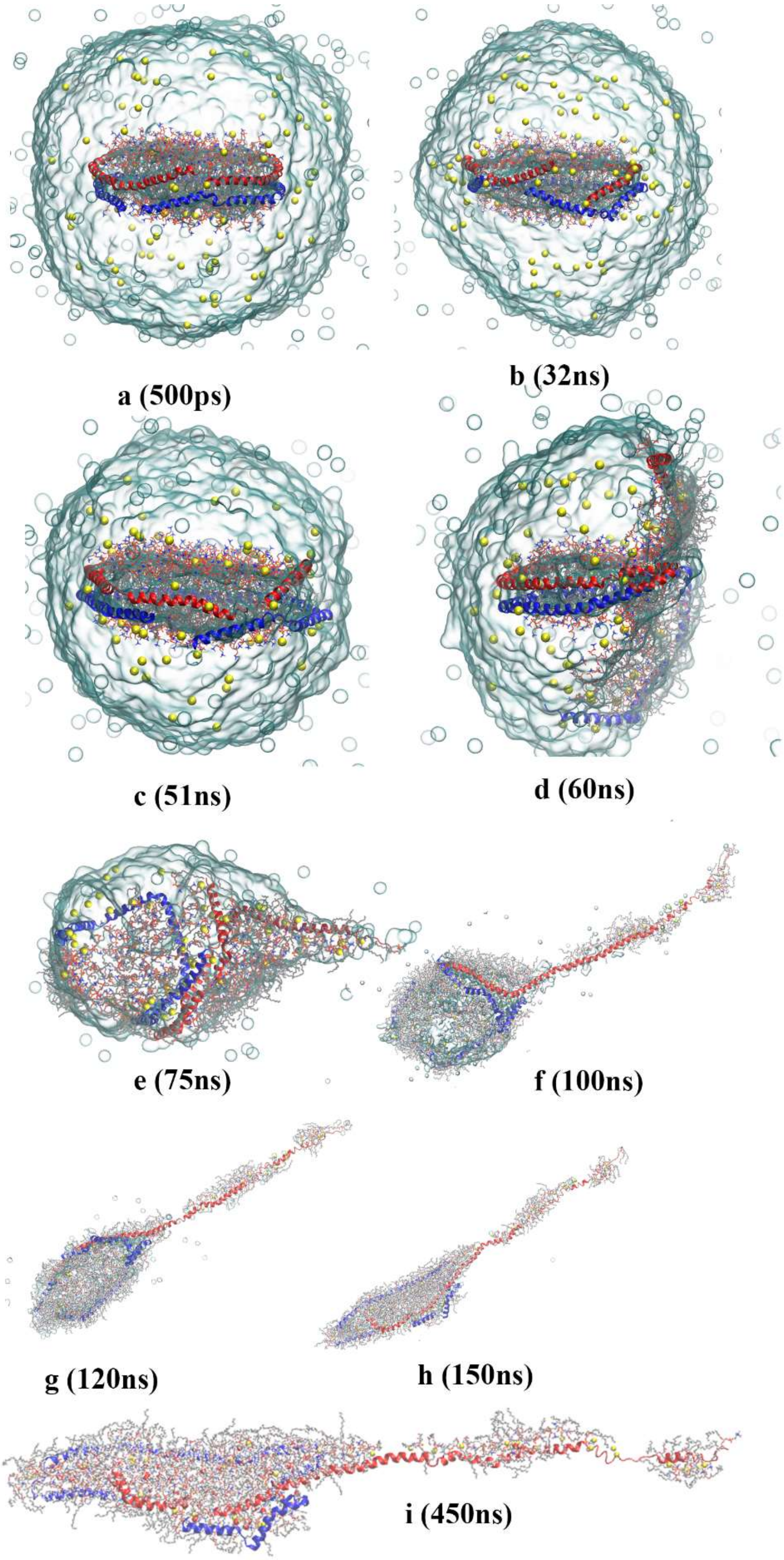


Figure S9. Evolution of the charged nanodisc droplet in the 370-2 simulation. The representation is the same as Figure 1.

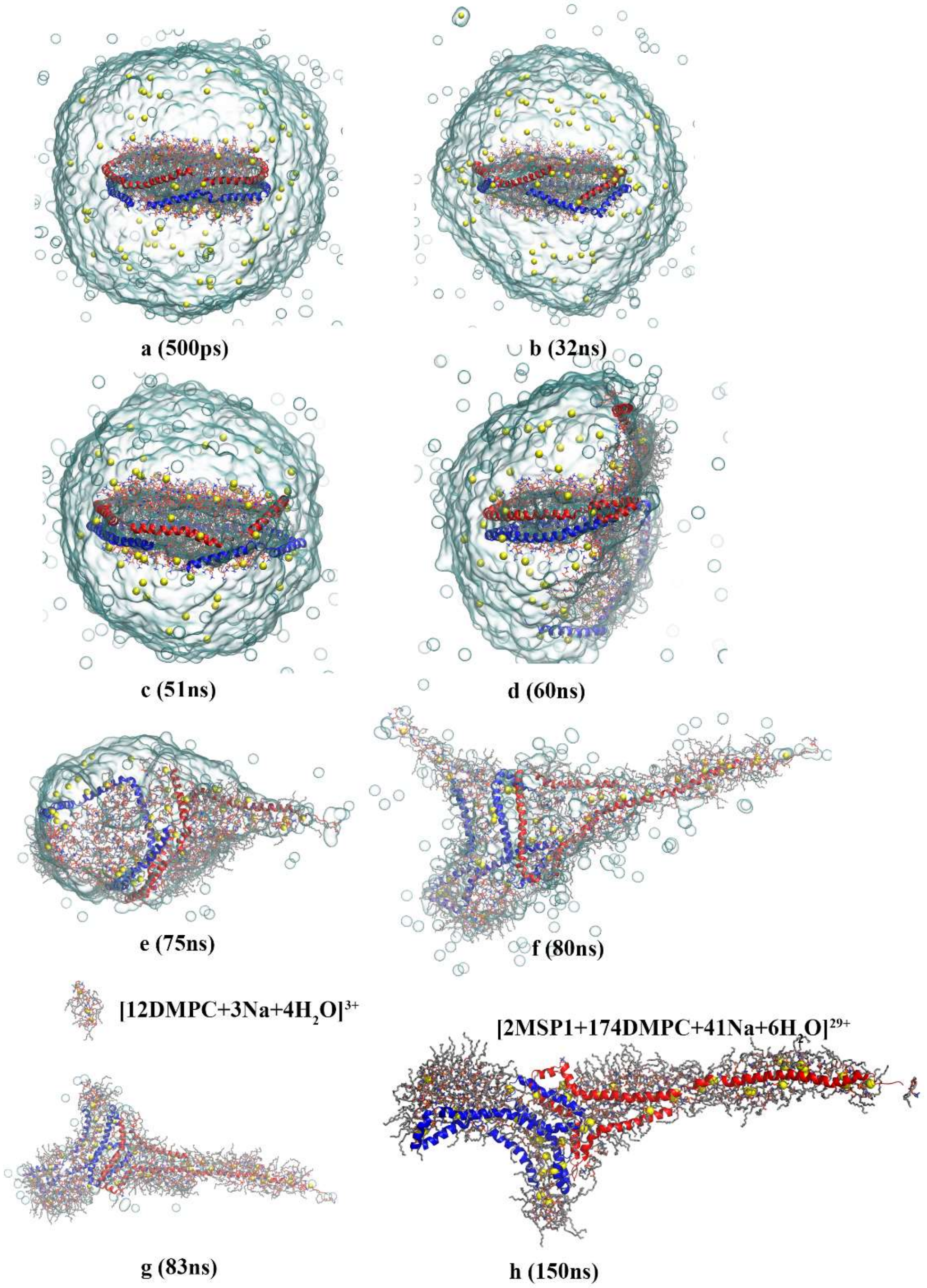


Figure S10. Evolution of the charged nanodisc droplet in the 450-2 simulation. The representation is the same as Figure 1.

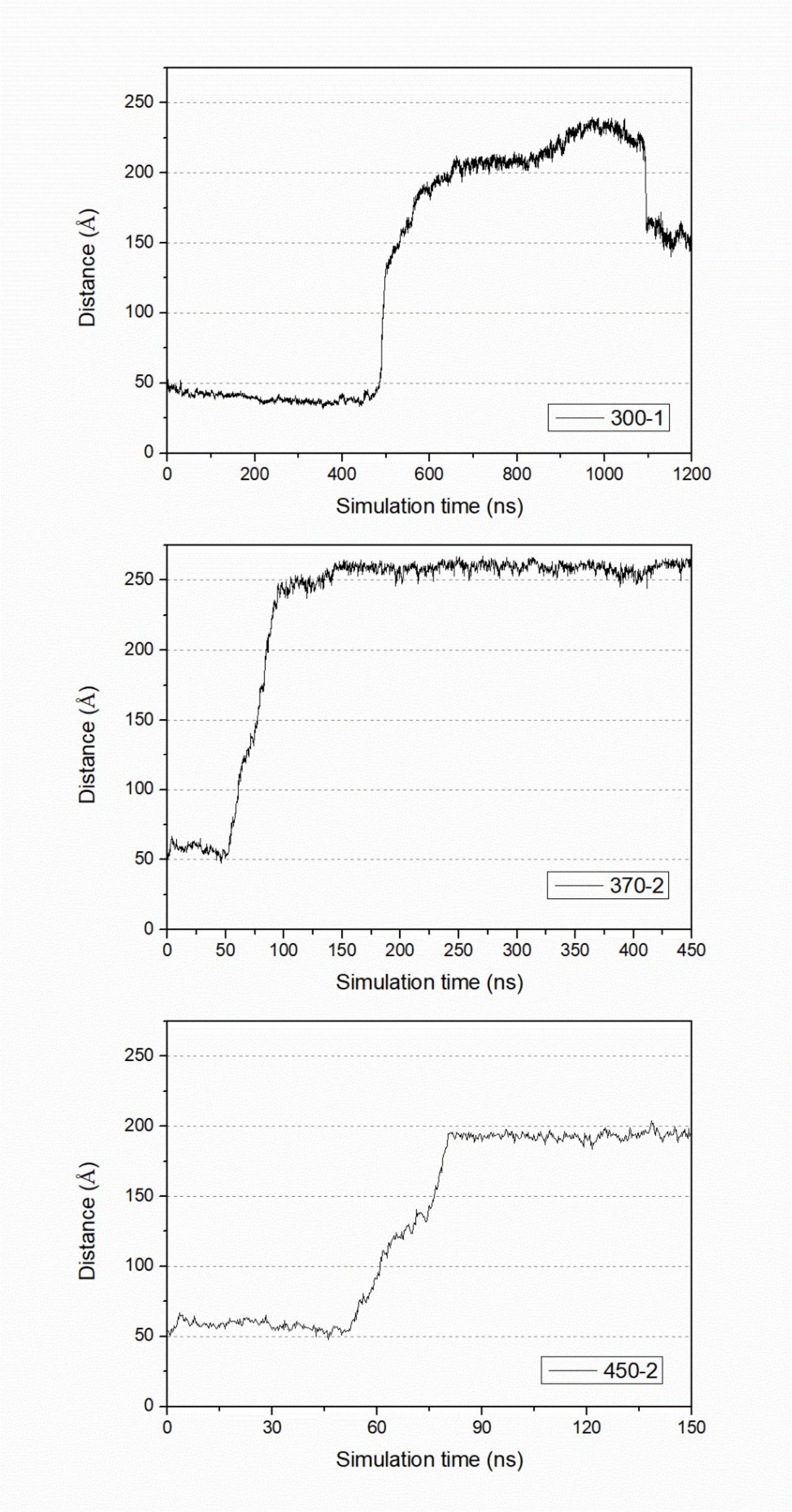


Figure S11. The variation of distance between the expeled teminal and the center of mass of the other monomer along simulation times in the three off-center trajectories (300-1, 370-2 and 450-2).

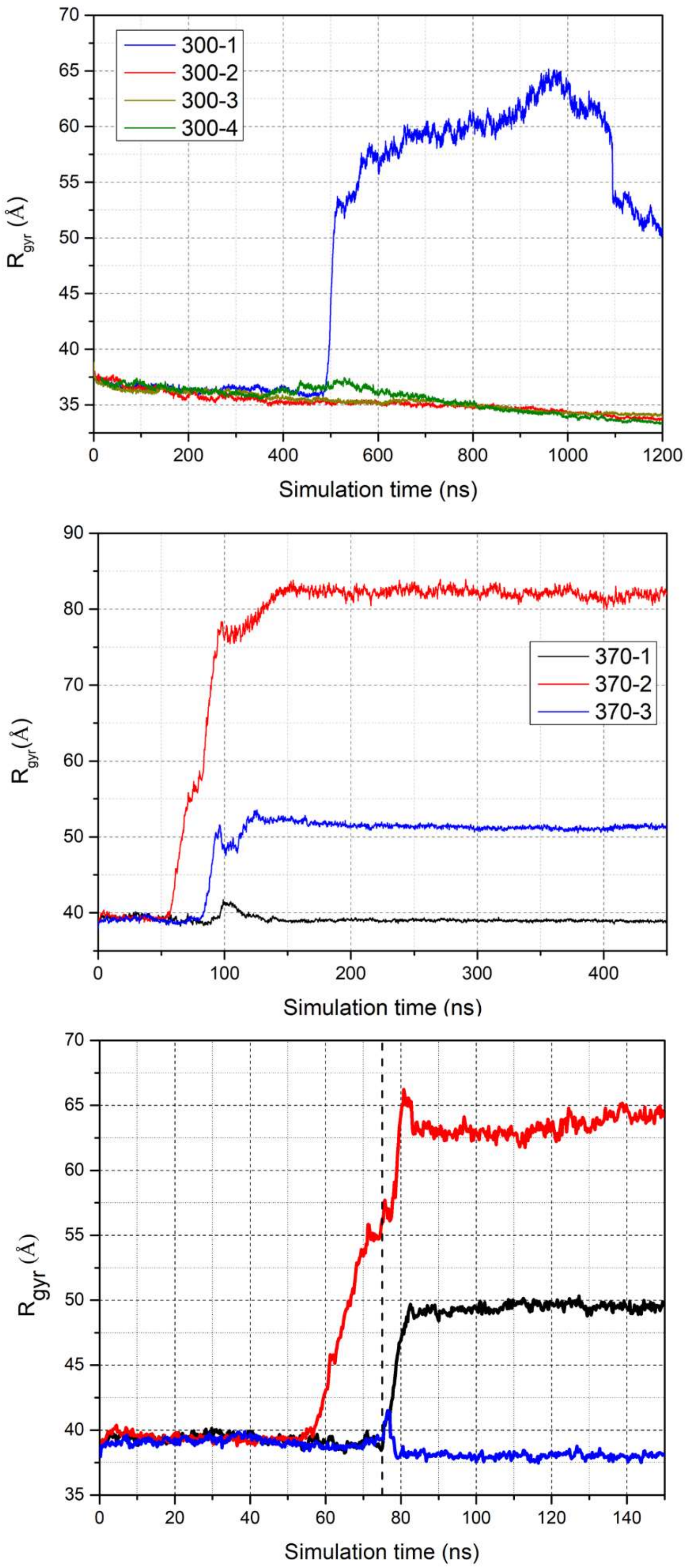


Figure S12. The evolutions of radius of gyration ($R_{gyr}$) of the nanodisc in all simulations at 300K, 370K and 370→450K from top to bottom.

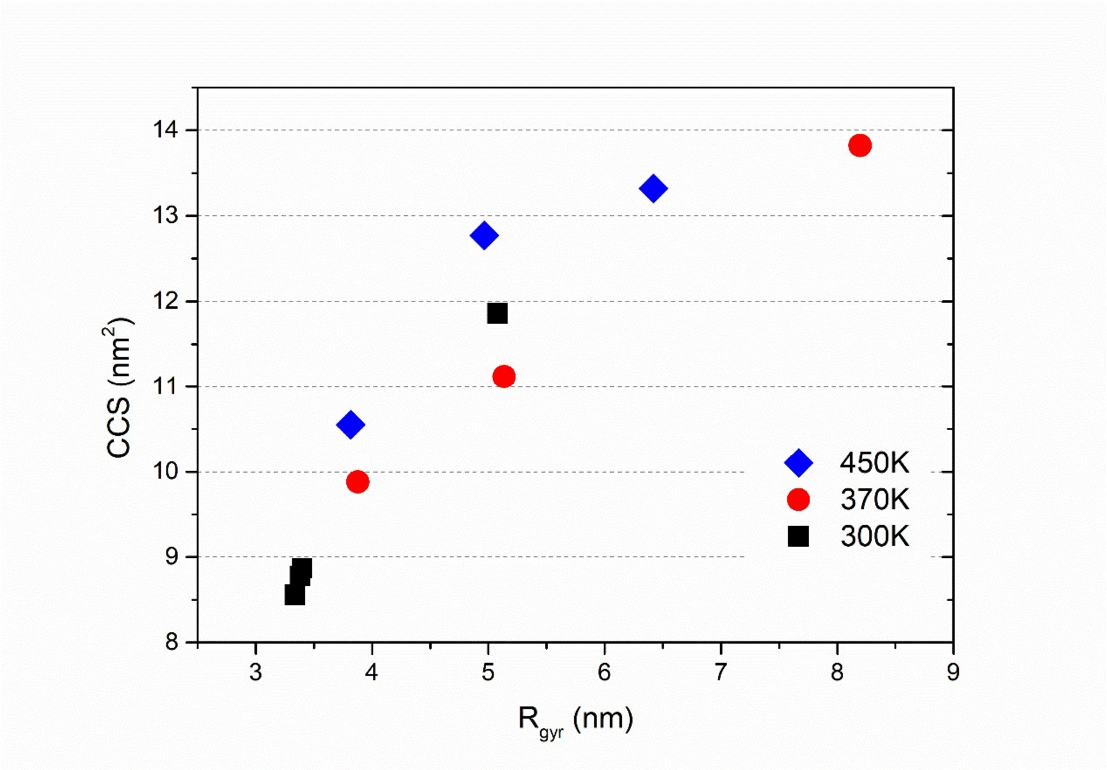


Figure S13. The correlation between the radius of gyration ($R_{gyr}$) and the collision cross section (CCS).

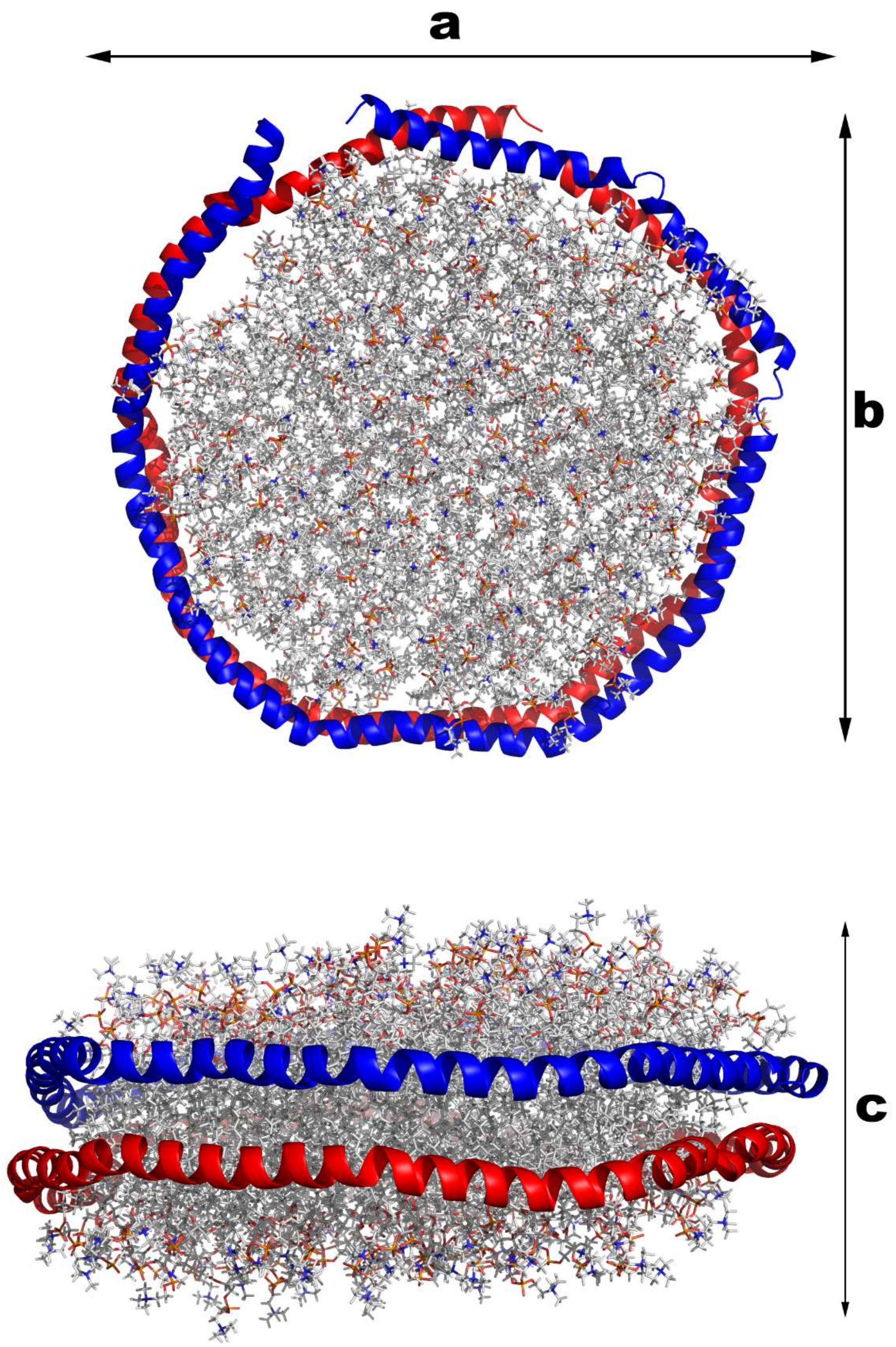


Figure S14. The three principal axes of the inertia tensor: a for the longest axis, b for the middle axis and c for the shortest axis.

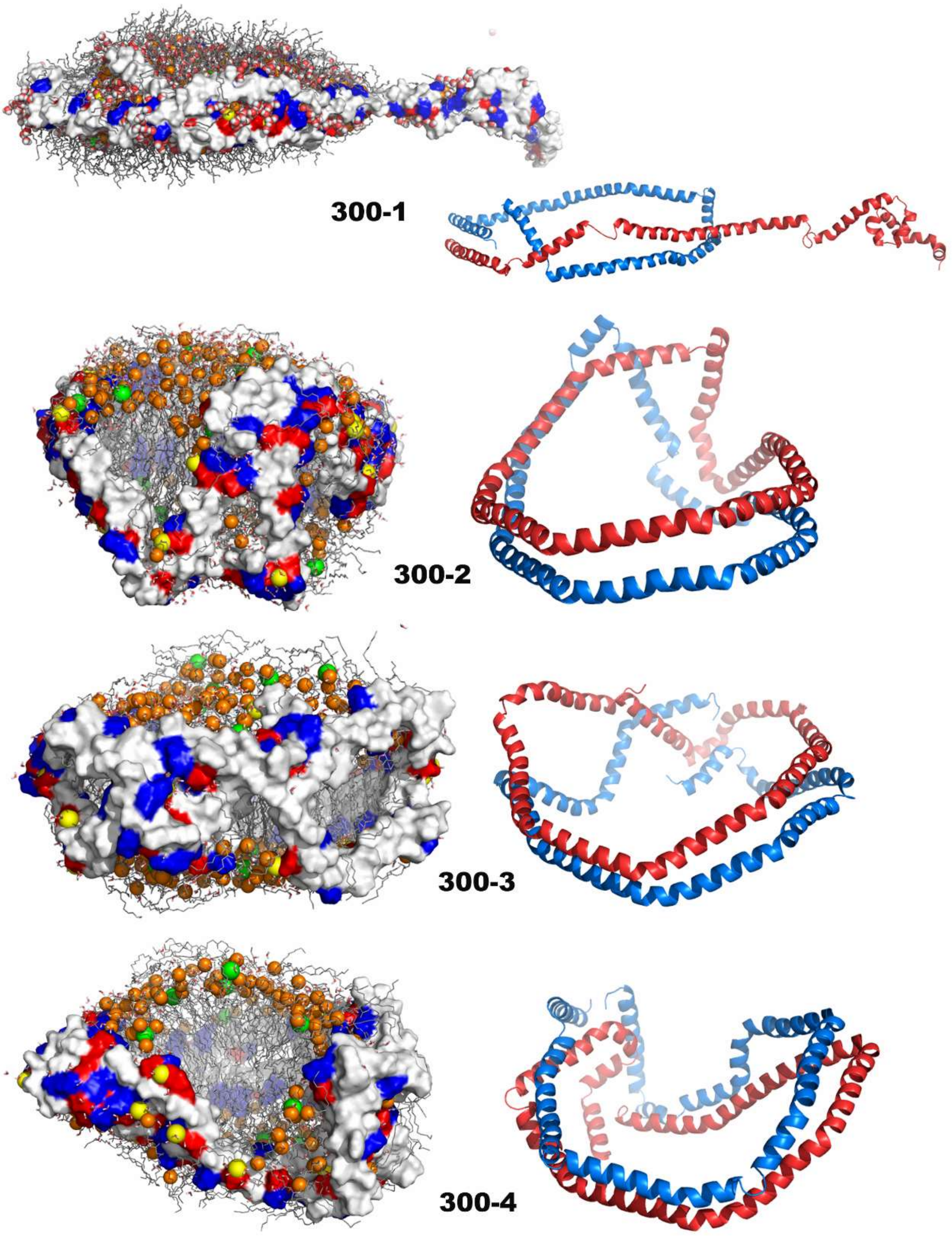


Figure S15. The structures of product gaseous ions of the simulations at 300 K and their MSP conformations. The representation is the same as Figure 3.

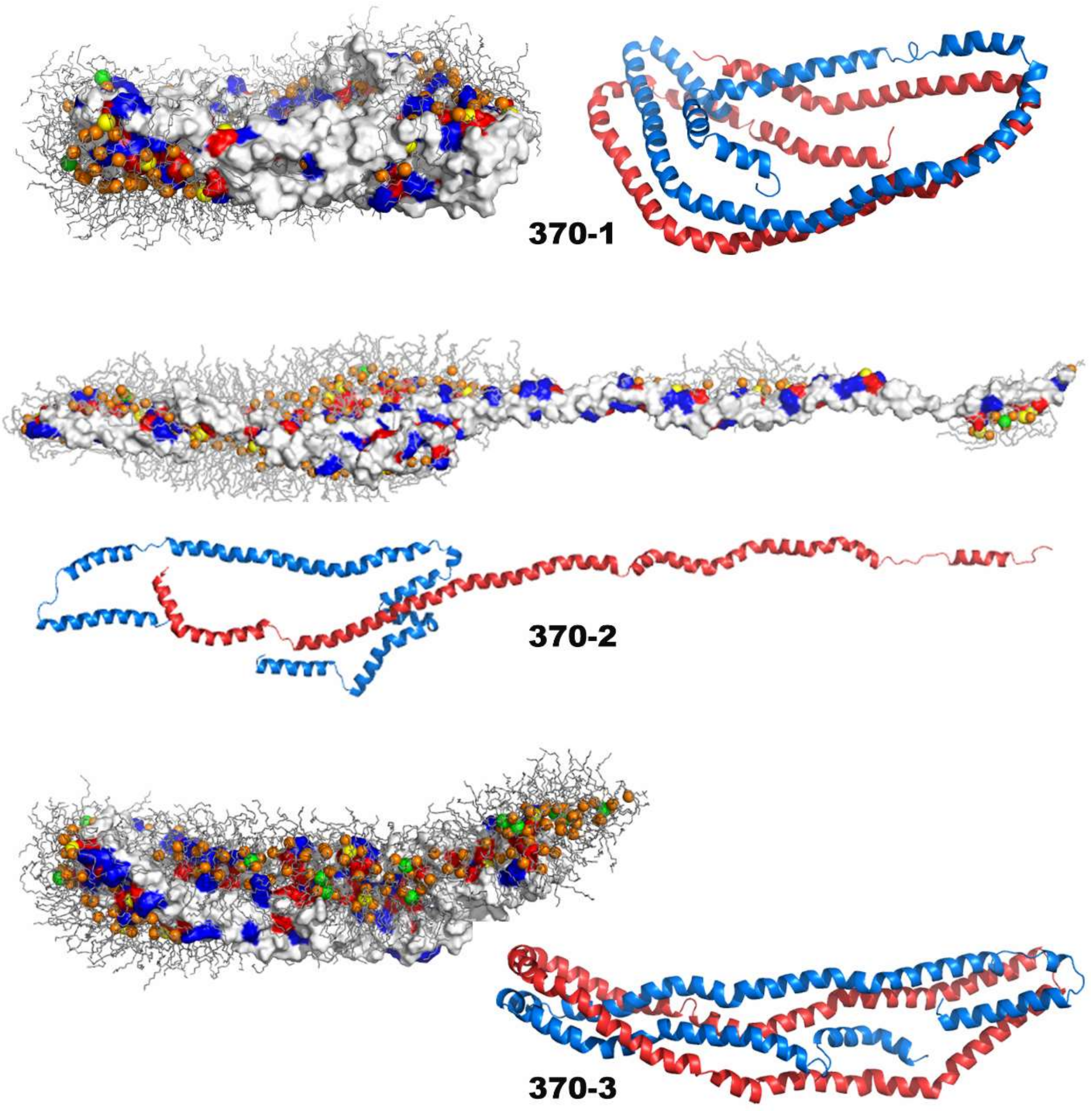


Figure S16. The structures of product gaseous ions of the simulations at 370 K and their MSP conformations.

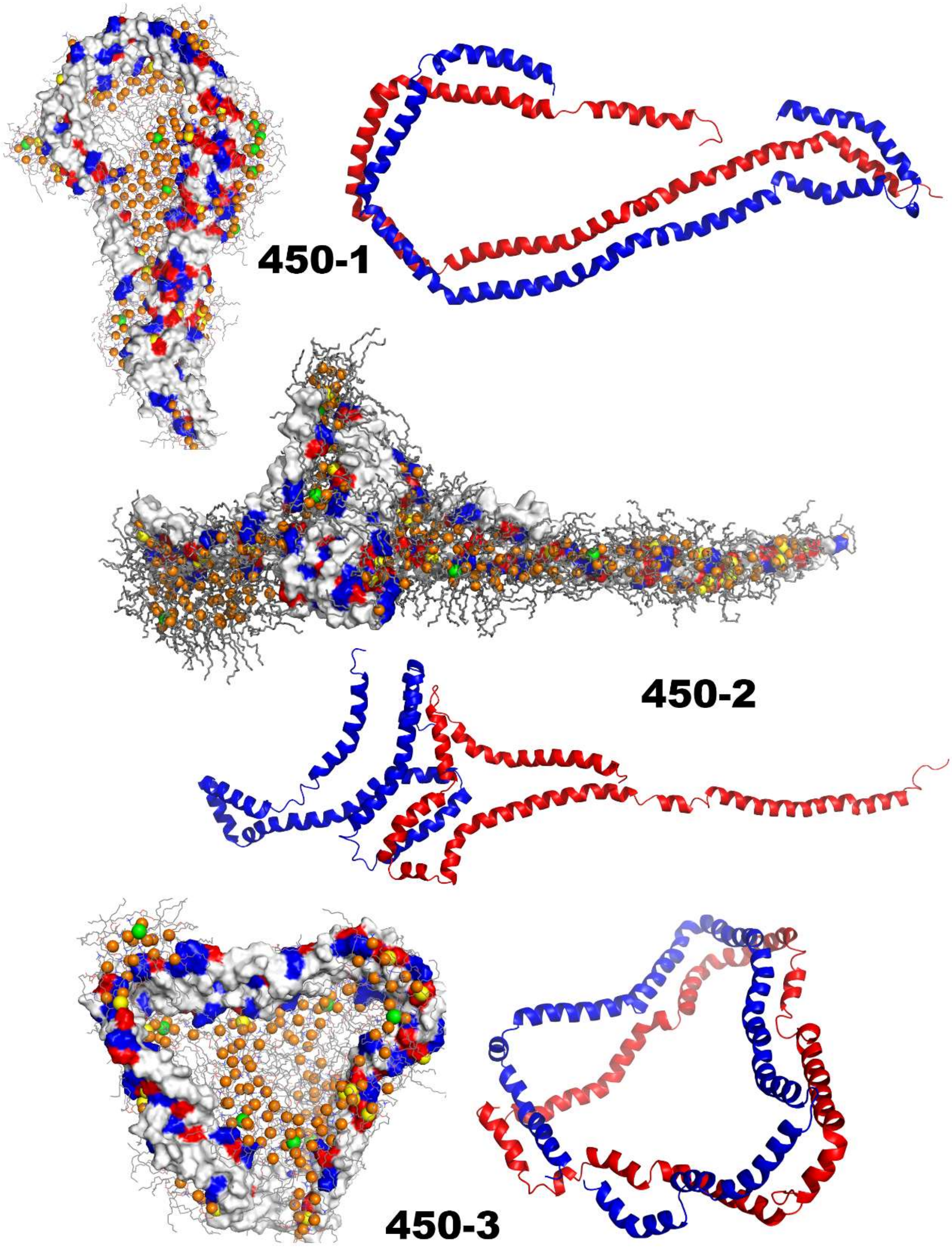


Figure S17. The structures of product gaseous ions of the simulations at 370→450 K and their MSP conformations.

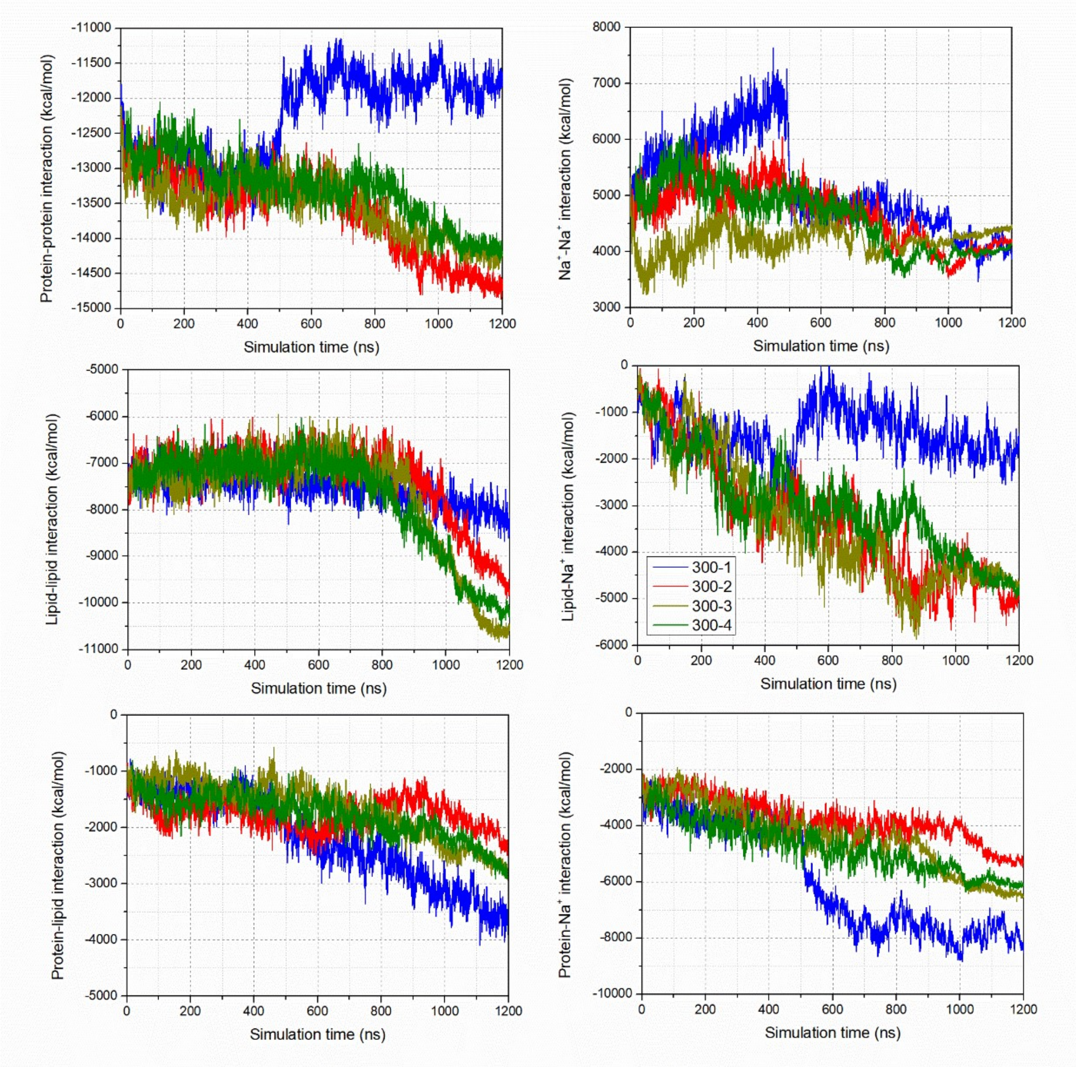


Figure S18. The evolution of electrostatic interactions between different components in the droplets in four simulations under 300K (300-1 in blue, 300-2 in red, 300-3 in dark yellow and 300-4 in green).

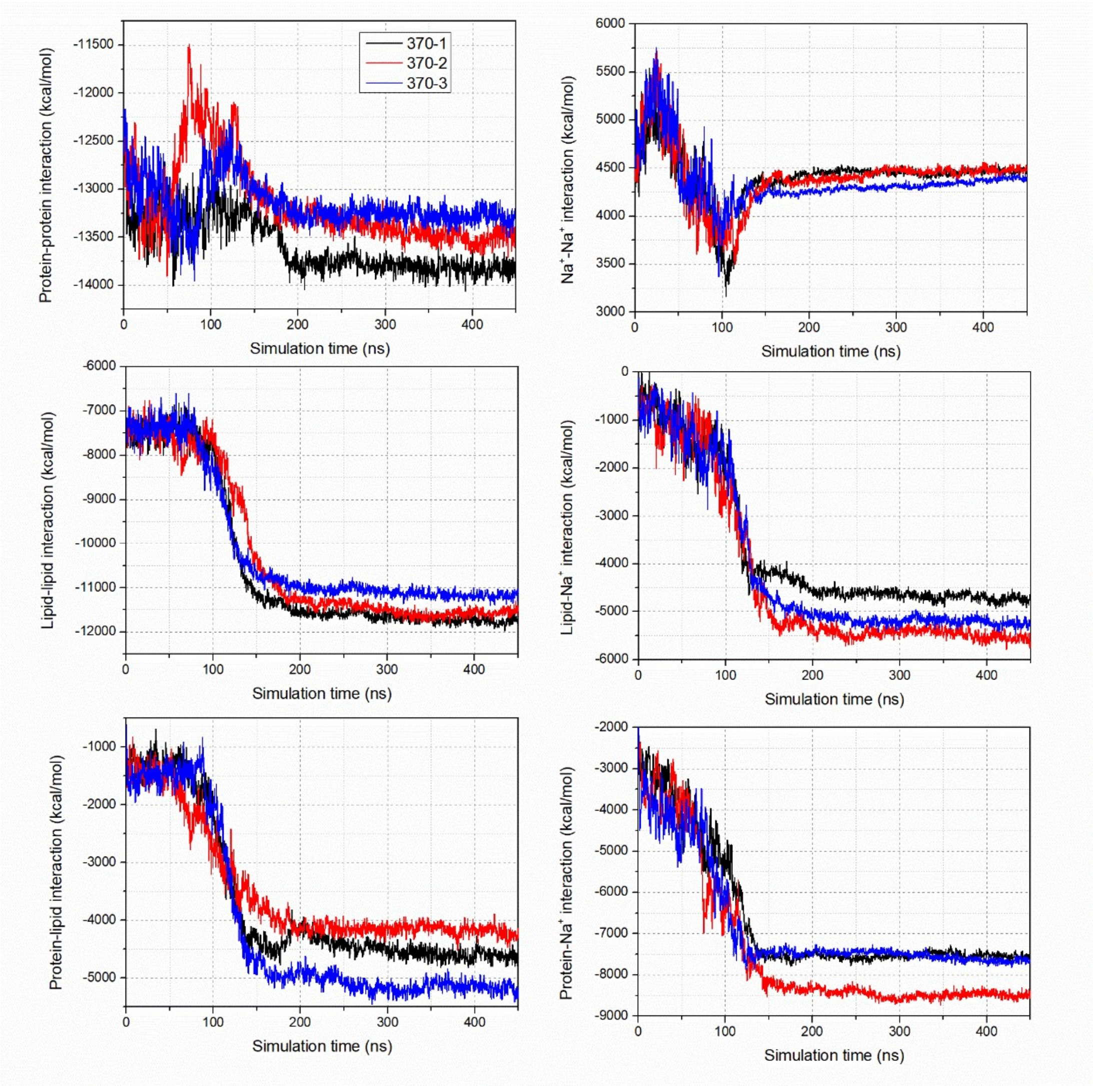


Figure S19. The evolution of electrostatic interactions between different components in the droplets in three simulations under 370K (370-1 in black, 370-2 in red and 370-3 in blue).

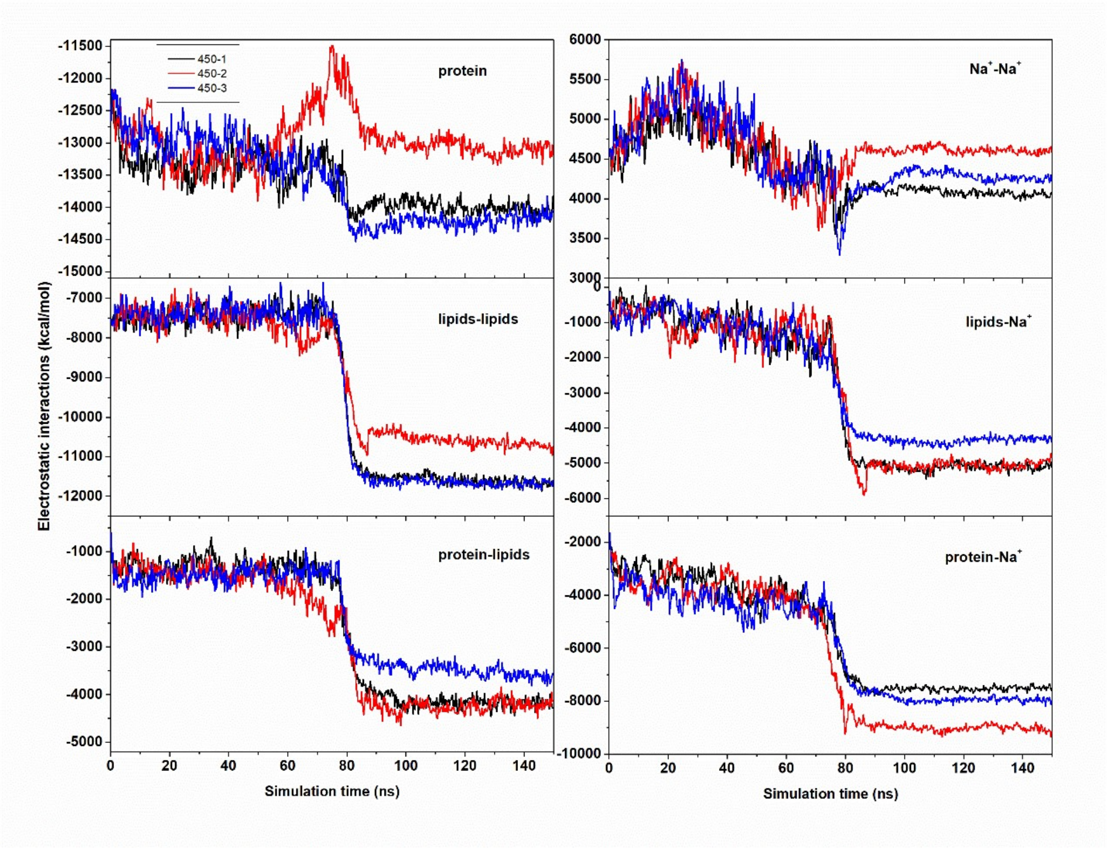


Figure S20. The evolution of electrostatic interactions between different components in the droplets in three simulations with 370→450K (450-1 in black, 450-2 in red and 450-3 in blue).

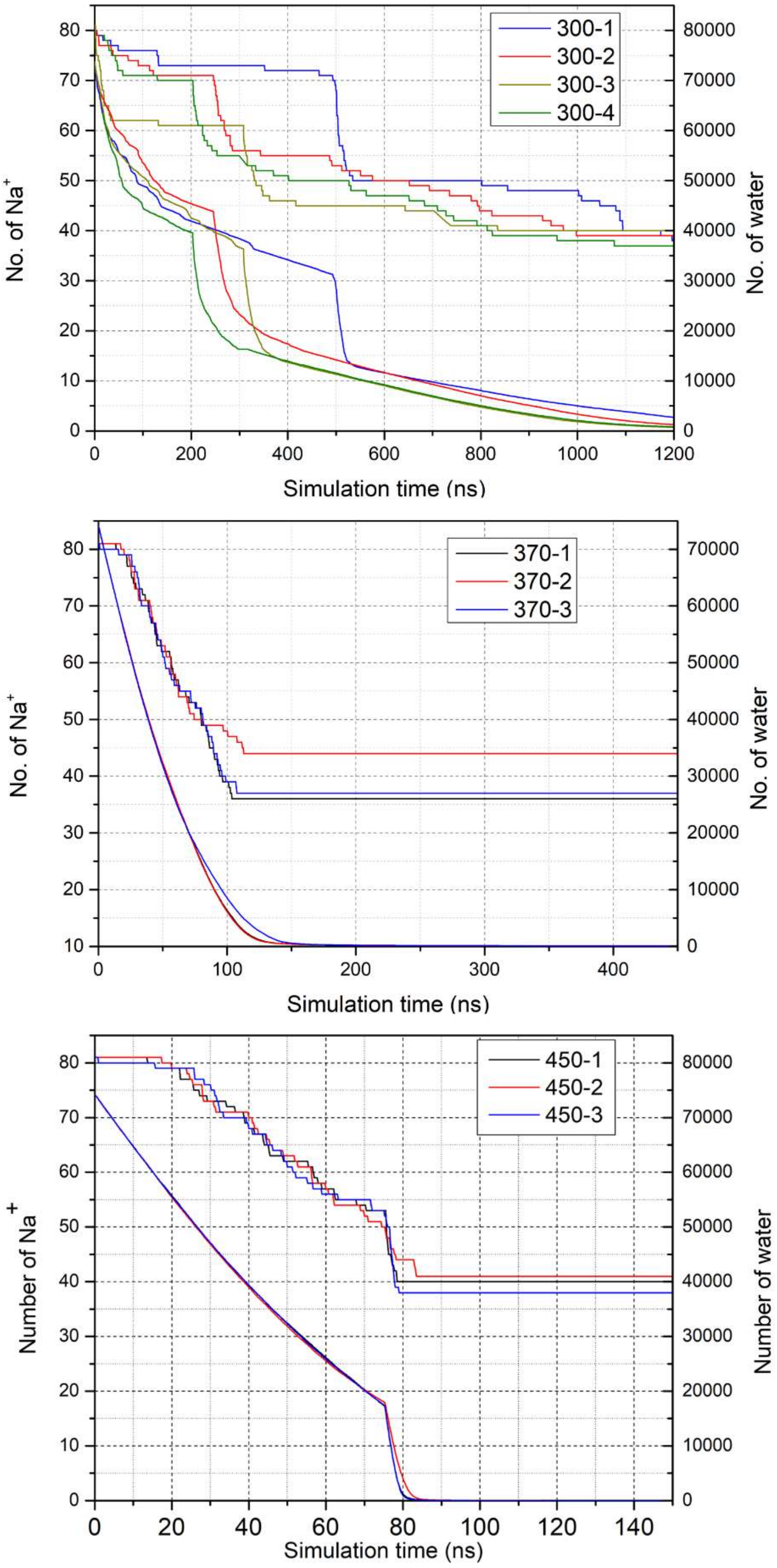


Figure S21. The evolution of the number of water and $Na^+$ molecules in remaining nano-droplets in all simulations. The simulations, 300-1, 370-2 and 450-2, follow the off-center process, and the remaining seven simulations follow the at-center process.

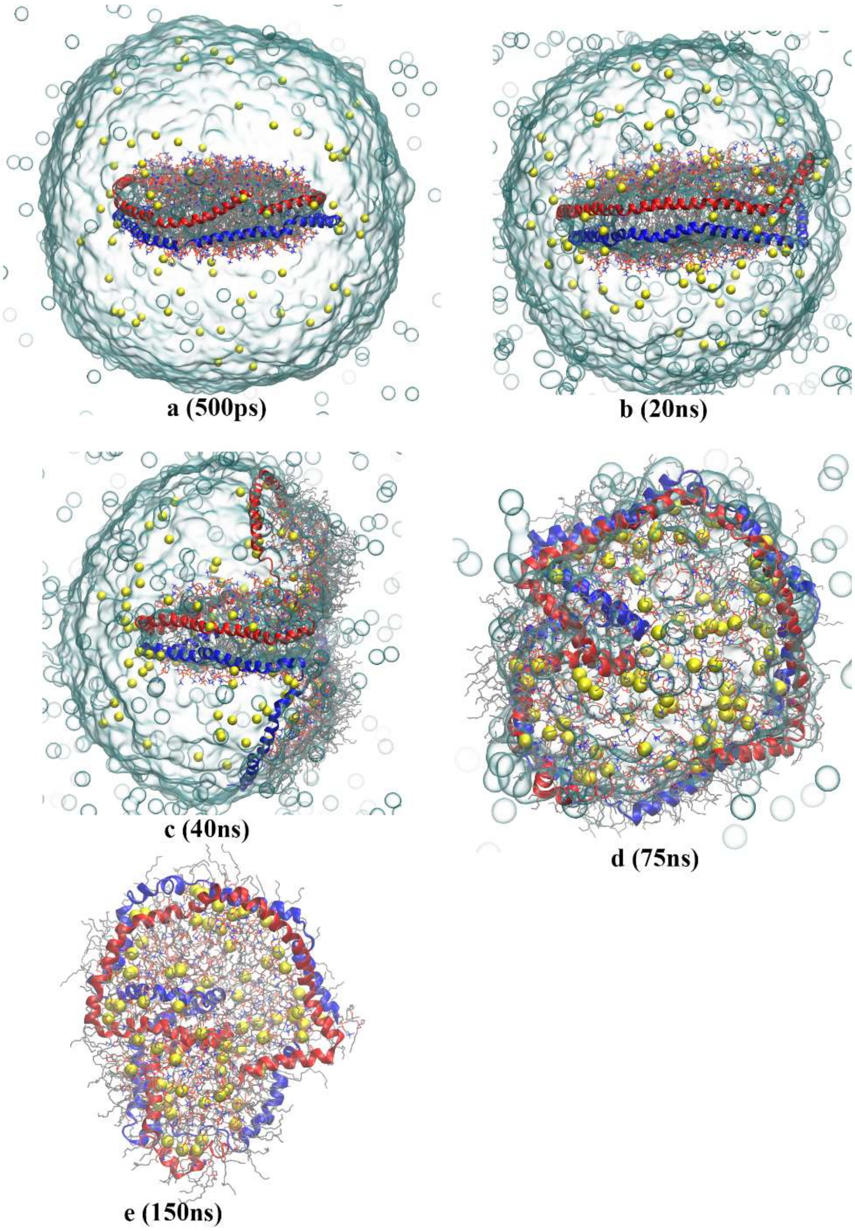


Figure S22. Evolution of the nanodisc droplet in the C12 simulation. The representation is the same as Figure 1. The composition of the gaseous ion is [2MSP1+186DMPC+81Na+3$H_2O$]$^{69+}$.

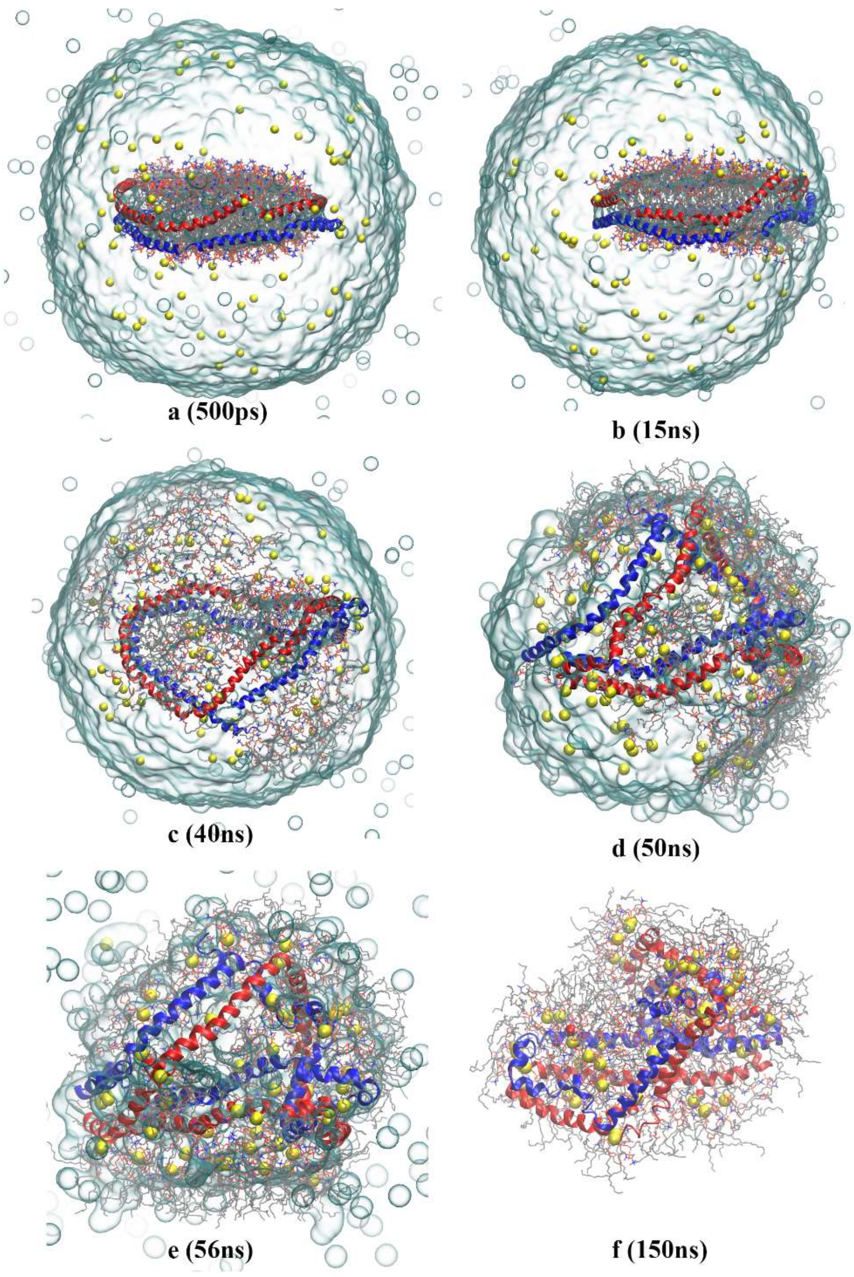


Figure S23. Evolution of the nanodisc droplet in the C33 simulation. The representation is the same as Figure 1. The composition of the gaseous ion is $[2MSP1+186DMPC+54Na+12H_2O]^{42+}$.

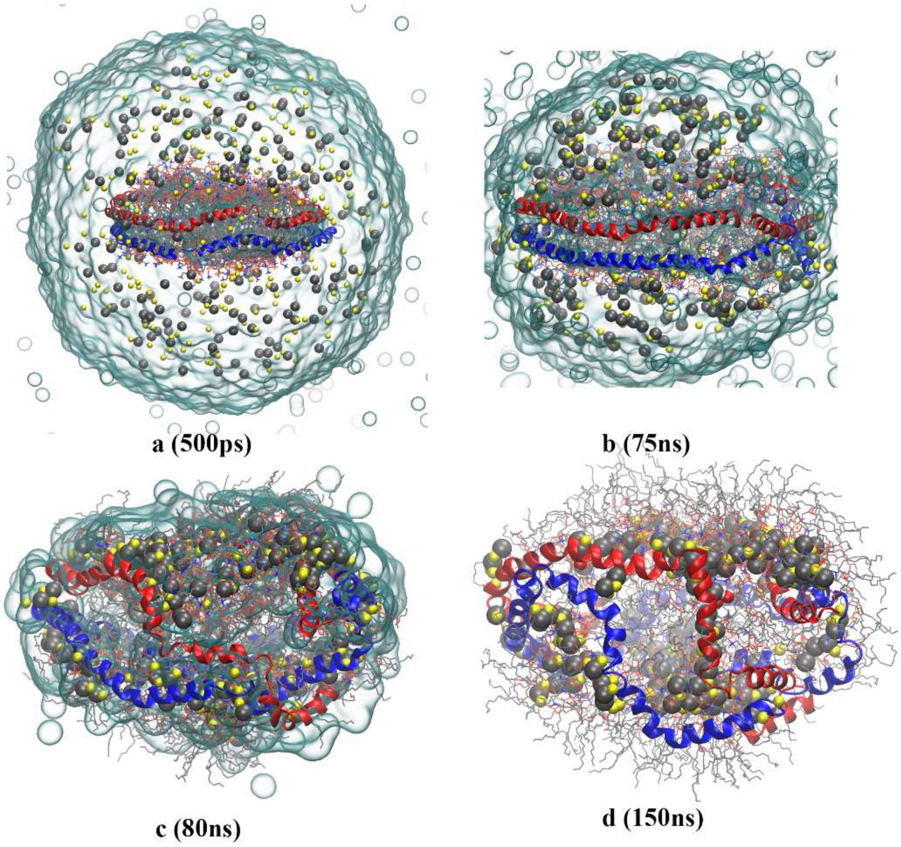


Figure S24. Evolution of the nanodisc droplet in the Neutral simulation. The representation is the same as Figure 1. In addition, the $Cl^-$ ions are shown in tan spheres. The composition of the gaseous ion is $[2MSP1+186DMPC+196Na+183Cl+4H_2O]^{1+}$.

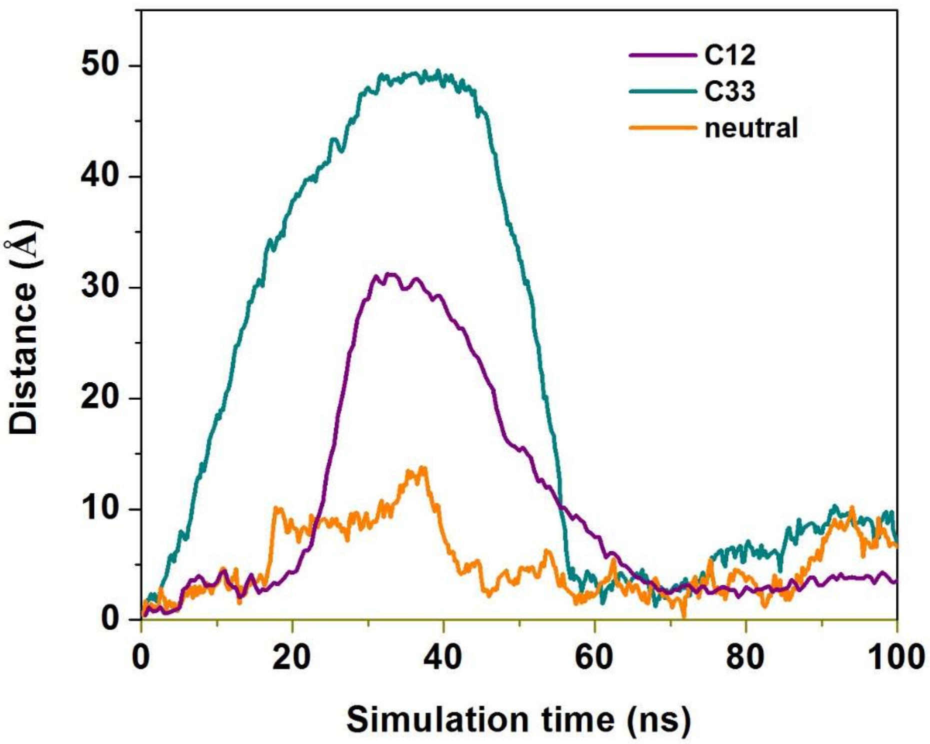


Figure S25. The distance between centers of mass of water molecules and nanodisc in the simulations, C12, C33 and Neutral (Table S1).

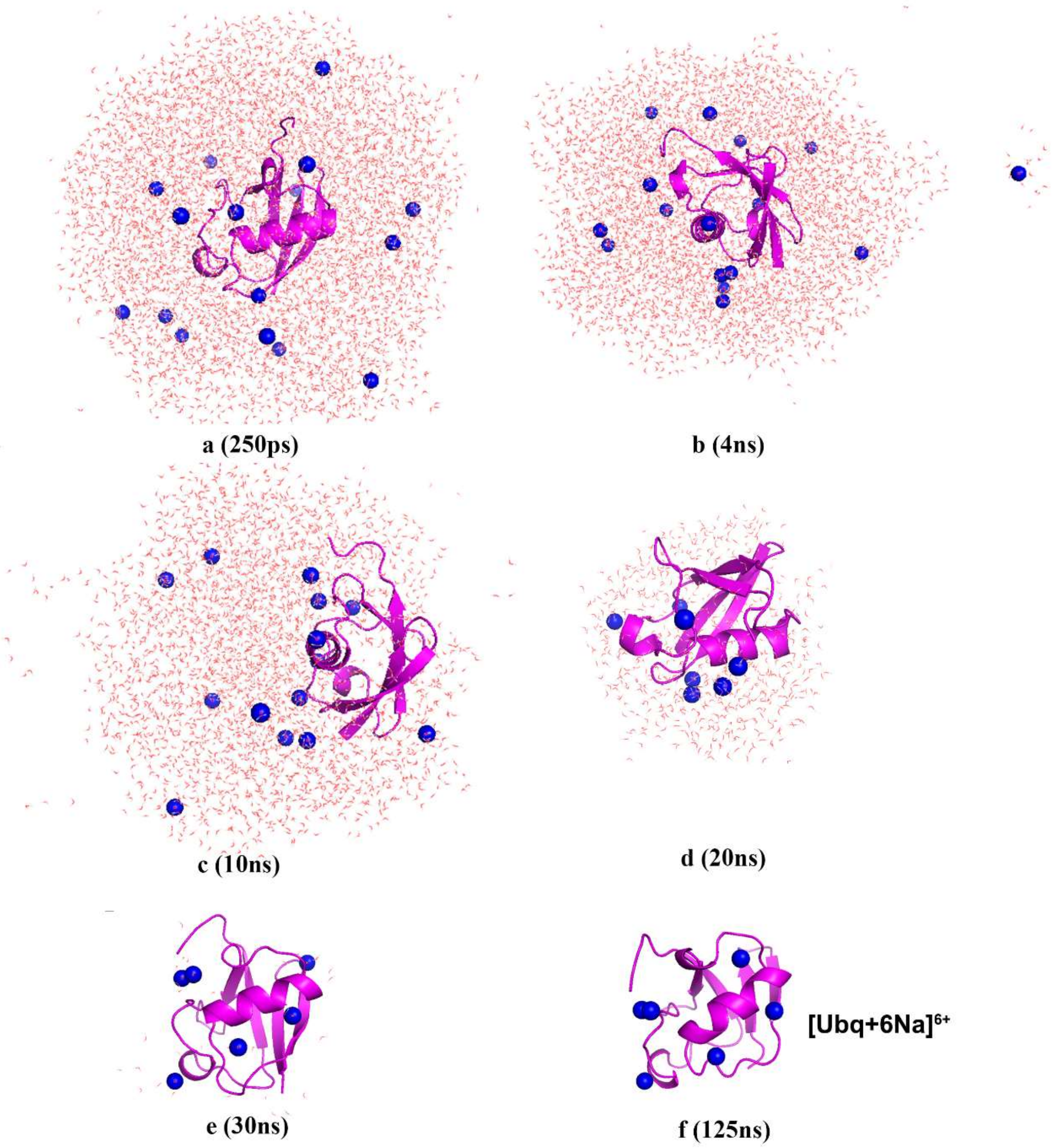


Figure S26. Evolution of the droplet with a net charge of 16+ in the ESI simulations of Ubq. The protein is shown in magenta cartoon, $Na^+$ in blue spheres.
